# Doxorubicin-induced Cardiotoxicity is Propagated by Paracrine Signaling through Small Extracellular Vesicles

**DOI:** 10.64898/2026.02.11.705398

**Authors:** George Ronan, Lara Ece Celebi, Nicole Kowalczyk, Noor Behnam, Jun Yang, Lauren Hawthorne, Frank Ketchum, Aoife J. Lowery, Michael J. Kerin, Pinar Zorlutuna

## Abstract

Cardiovascular disease (CVD) is the leading cause of death in the United States and worldwide. While most of these deaths are the result of chronic heart diseases, some CVDs are induced artificially. Doxorubicin (DOX) is a chemotherapeutic that is commonly used to treat breast cancer which is one of the most common types of cancer in the United States. While DOX is an effective anti-cancer agent, over 10% of treated women show signs of acute cardiotoxicity immediately following treatment, and approximately 2% develop severe cardiotoxicity up to 10 years after the end of treatment. Despite this prevalence, the mechanism by which the onset of this cardiotoxicity occurs over time is not well understood. Here, we show that treatment of cardiac cells with DOX changes the cardiac function and the resulting paracrine signaling profile. Subsequent exposure of healthy cells to these altered paracrine agents can recapitulate the effects of direct DOX exposure in 2D and 3D *in vitro* models. We suggest that this is the result of an altered paracrine miRNA profile and other paracrine factors that propagate the initial disruption caused by direct DOX exposure. Plasma EV miRNA profiling of blinded patient samples revealed distinct clustering by DOX-cardiotoxicity risk, with high-risk patients exhibiting miRNA signatures similar to those from DOX-treated tissue-engineered models. Pathway analysis of the most distinguishing miRNAs linked them to cardiac homeostasis and cardiotoxicity-related mechanisms, supporting the potential of plasma EV miRNAs as noninvasive biomarkers for early risk stratification and personalized cardioprotective interventions in oncological care, and the targeting of key clusters of miRNAs to enhance both understanding of and intervention strategies for preventing the onset of DOX cardiotoxicity.

## Introduction

Cardiovascular disease (CVD) is the leading cause of death in the United States and worldwide^1^. While most of these deaths are the result of chronic heart diseases which lead to heart failure after myocardial infarction (MI), some CVDs can be induced artificially, such as via off-target effects of other therapies^2^. This is particularly common in chemotherapy and remains a pervasive issue with both established and novel targeted chemotherapeutics^3^, often being referred to as “off-target toxicity”. Off-target effects which compromise or otherwise damage cardiac health and functionality are commonly referred to as “off-target cardiotoxicity”, although there is little consensus on a more specific definition for this term in literature^4^. For many chemotherapeutics, the cause of cardiotoxicity can be linked directly to exposure to the chemotherapy agent^5^, spurring advances in chemotherapy delivery vehicles and targeting techniques^3,4^. For some chemotherapeutics, however, the mechanism by which cardiotoxicity occurs is less clear.

Doxorubicin (DOX), an anthracycline, is a chemotherapeutic that has been commonly used to treat breast cancer, one of the most common types of cancer in the United States. Breast cancer affects 1 in 8 women both in the United States and globally and comprises 30% of yearly cancer diagnoses and 12.5% of cancer diagnoses globally^6^. Since the initial formulation of DOX in the 1960’s and approval for medical use in 1974, DOX has proven to be highly effective in treating cancers including breast, bladder, stomach, lungs, ovarian, thyroid, soft tissue sarcoma, multiple myeloma, lymphoma, and leukemia^7,8^, as have the more than 2000 DOX analogues. While DOX and anthracycline cocktails are very successful in mitigating or otherwise destroying breast cancer and other cancers, over 10% of women showed signs of acute cardiotoxicity immediately following treatment^7^ and approximately 2% developed severe cardiotoxicity up to 10 years after the end of treatment^9^, despite known clearance of DOX in less than 48 hours^10^. This DOX-induced cardiotoxicity has since been well-established in DOX and many of the DOX analogues, both alone and in chemotherapy cocktails, and is thought to primarily operate through transcriptional and mitochondrial damage^8,9^. However, the precise mechanisms by which DOX cardiotoxicity is initiated as well as how such effects could persist even after DOX clearance are the subject of intense debate^7^. We hypothesize that early DOX exposure pathologically disrupts the paracrine signaling of myocardial cells, which, over time, propagate this dysfunction to the surrounding tissue to eventually result in observable cardiotoxic effects. This can be demonstrated by independently recapitulating the effects of DOX treatment using only extracellular vesicles (EVs) isolated from DOX-treated cells.

Extracellular vesicles (EVs) are traditionally defined as apoptotic bodies (∼1 µm – ∼5 µm diameter), microvesicles (∼200 nm – ∼1000 nm diameter), and exosomes (∼30 nm – ∼200 nm diameter)^11^, with exosomes being particles of particular interest due to exosomes acting as major vehicles for paracrine and endocrine transfer of proteins and nucleic acids^12^. Recently, however, the identification of numerous difficult to separate subgroups of exosomes and non-exosome EVs with sub-200 µm diameter has developed into separate classification of EVs as medium or large EVs (mEVs or lEVs, respectively) with diameters typically greater than 200 µm, and small EVs (sEVs) with diameters typically less than 200 µm^13^. These sEVs, like exosomes, are commonly vehicles for the transport of nucleic acids and proteins, notably micro RNAs (miRNAs) and cytokines, which influence many diverse and pathologically relevant biological processes, including angiogenesis, immunomodulation, mitochondrial activity, and epithelial to mesenchymal transition^11,13^. Additionally, exosome-like sEVs have been demonstrated to be actors in chronic CVDs^14,15^, and have been suggested as actors in the onset of DOX-related cardiotoxicity^16^. While evidence suggests that paracrine signaling plays some role in the onset of DOX-cardiotoxicity, the proposed mechanisms and involvement in the propagation of cardiotoxicity are often confounding^16,17^. Some studies have suggested that sEVs directly maintain and transport DOX to mediate cardiotoxic effects, but current literature does not wholly support this and instead individual miRNAs, transported by sEVs, are under investigation as major actors^18,19^. We suggest that DOX exposure in cardiac cells induces a global shift in the miRNA population carried by sEVs, where the collective dysregulation of multiple miRNAs, not any single miRNA, drives the propagation of cardiotoxic effects. By characterizing a total miRNA population shift, rather than identifying a single miRNA, we are also able to identify novel miRNA biomarkers for DOX-related cardiotoxicity and validate them in clinical samples.

In this study, we show for the first time in literature that DOX-related cardiotoxicity can be propagated by paracrine factors independent of direct exposure to DOX in 2D, 3D, and heart-on-a-chip models. Furthermore, we identify novel biomarkers of DOX-related cardiotoxicity from sEVs in conditioned media of DOX-exposed cells and validate those markers in clinical samples. Finally, we perform downstream analysis of identified miRNA targets to ascertain potential involved pathways for future therapeutic intervention, and cross reference with miRNAs and pathways currently suspected of involvement in DOX-related cardiotoxicity. The identification of sEVs as crucial elements in the onset and propagation of DOX-related cardiotoxicity will help elucidate specific mechanisms of both acute and chronic cardiotoxicity, as well as assist in the development of enhanced chemotherapeutic approaches to limit off-target cardiotoxic effects.

## RESULTS

### DOX-EVs Induce Similar Dysfunction to Direct DOX Treatment in Cardiac Muscle and Stromal Cells

To assess the effects of DOX-EVs on cardiac cells compared to direct DOX treatment, hiPSC-derived cardiomyocytes (iCM) and cardiac fibroblasts (iCFs) were treated with a blank control, DOX-conditioned media, or media conditioned with EVs from DOX-treated iCFs (Figure 1A). EVs from iCFs were used in 2D culture, as CFs act as master regulators of the myocardial microenvironment and local signaling *in vivo*^20^.

**Figure 1:**
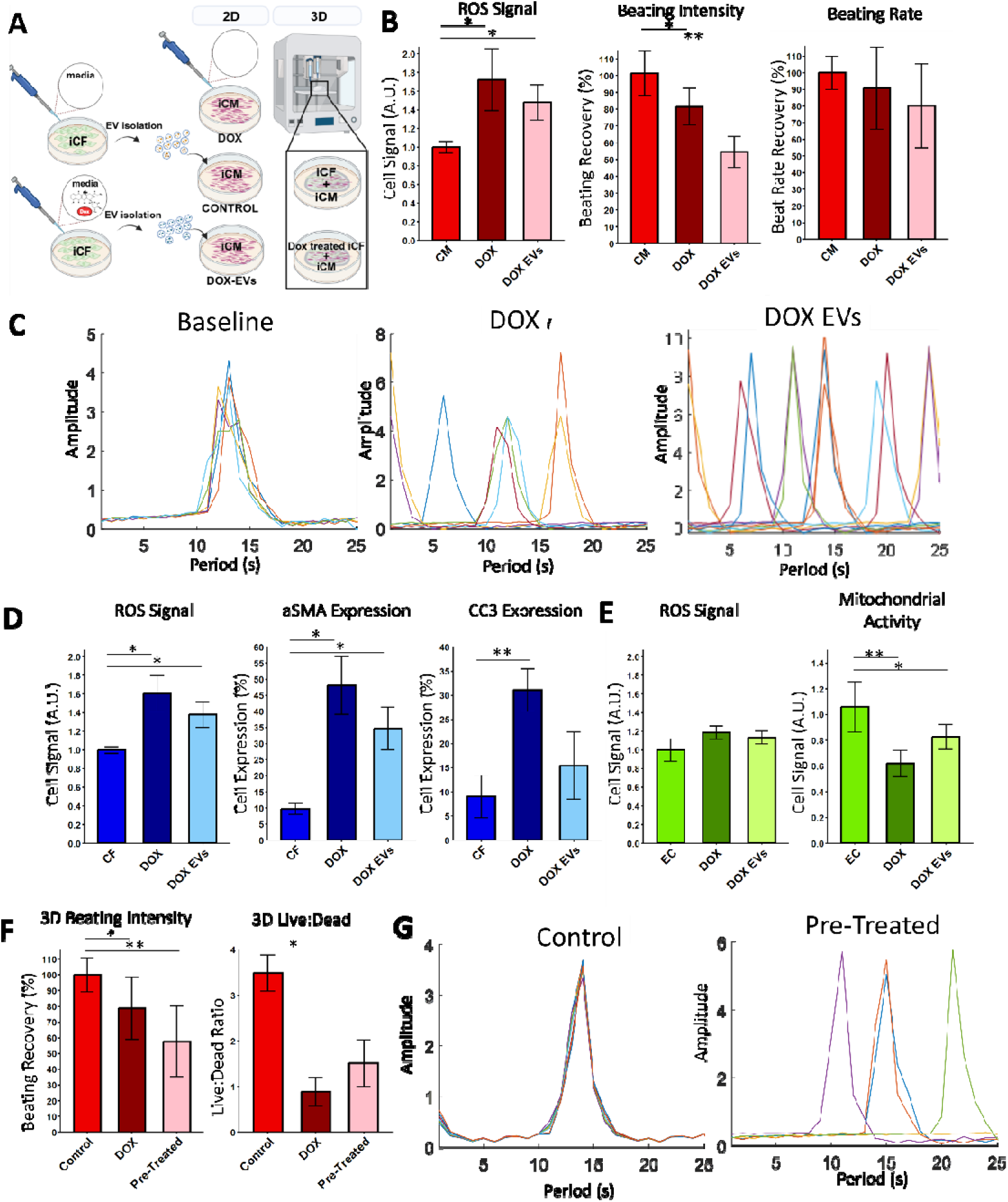
EVs from DOX-Treated Cells can Independently Recapitulate DOX Toxicity in vitro. (A) Experimental setup for assessing DOX-induced cardiotoxicity in 2D and 3D culture systems. (B) ROS level (left), beating intensity (middle), and beating rate (right) of iCMs when treated with DOX, DOX-EVs, or a PBS blank for 48 h. (C) Overlaid beating patterns of iCMs by cohort. (D) ROS level (left), αSMA expression (middle) and cleaved caspase-3 expression (right) of iCFs by cohort. (E) ROS level (left) and cell metabolic activity (right) of iECs by cohort. (E) Beating intensity (left) and cell survival (right) 48 h after seeding with control or DOX-treated iCFs with iCMs in a 3D model, and treated with control or DOX media. (F) Overlaid beating patterns of 3D models with control (left) or pre-treated (right) iCFs. n ≥ 3 for all groups tested, * p < 0.05, ** p < 0.01, *** p < 0.005, **** p < 0.001 assessed by one-way ANOVA with Tukey’s post-hoc.

First, due to the known effects of DOX in inducing oxidative stress in cardiac cells^21^, relative ROS signal was assessed for each cohort. Interestingly, DOX-EV treatment significantly increased ROS signal in iCMs (32%), though not as much as direct DOX treatment (65%) (Figure 1B, left). Beating analysis of iCMs treated for 48 h with either the control, DOX media, or DOX-EV media (Supplemental Video 1-2-3) revealed that both treatment groups reduced beating velocity (DOX, 27%; EV, 54%) (Figure 1B, middle) and a modest decrease in average beating rate (DOX, 9%; EV, 20%) (Figure 1B, right).

To better assess changes in beating rate and overall beating regularity, a live Ca^2+^ stain was used to measure the beating behavior of the iCMs temporally (Supplemental Video 4-5-6). The control cells demonstrated a regular period of ∼4.5 s, where the DOX-treated cells demonstrated an irregular period ranging from ∼2 s to ∼6 s, and the DOX-EV group showed a similar period to DOX-treated cells, ranging from ∼2 s to ∼5 s (Supplemental Figure S1). When overlayed, the control peaks tended to overlap, while both the DOX and DOX-EV group peaks tended to appear sporadically (Figure 1C). Beating signals were transformed via fast Fourier transform (FFT) and decomposed into sine wave components to allow for more direct comparison of the signals. The control group showed regular peaks of decreasing amplitude, as expected, whereas both the DOX and DOX-EV groups showed irregular peaks with random amplitude (Supplemental Figure S2).

Next, the differences in the effects of DOX-EVs and direct DOX treatment on iCFs were evaluated. First, relative ROS signal was assessed for each cohort. As with iCMs, both DOX treatment and DOX-EV treatment significantly increased ROS signal (Figure 1D, left). Additionally, both DOX and DOX-EV treatment induced a significant increase in αSMA expression, with DOX increasing expression by nearly 300% compared to the control and DOX-EVs inducing a nearly 200% increase over the same time period (Figure 1D, middle). Additionally, DOX treatment resulted in a more than 3-fold increase in daily cell death rate, from ∼3% to over 15%, though this was not replicated by treatment with DOX-EVs alone (Figure 1D, right).

### DOX Treatment of iECs Induces Different Dysfunction from other Cardiac Cells

In addition, the effects of direct DOX treatment and DOX-EV treatment on hiPSC-derived endothelial cells (iECs) were evaluated. As with iCMs and iCFs, ROS signal was evaluated under all three treatment conditions. Interestingly, however, neither DOX nor DOX-EVs induced a significant increase in ROS expression in iECs (Figure 1E, left), though both DOX and DOX-EV groups showed a non-significant increase. To evaluate overall metabolic rate of iECs under each treatment, a cell metabolism assay was run on each cohort. In this case, both DOX and DOX-EV-treated iECs showed significantly decreased mitochondrial activity compared to the control (Figure 1E, right). Furthermore, the decrease observed in both DOX and DOX-EV groups was very similar.

### Pre-treatment of iCFs with DOX Induces DOX-like dysfunction in 3D Cardiac Models

To assess the ability of local signaling to propagate DOX-cardiotoxicity, 3D bioprinted co-culture models of iCMs and iCFs were constructed. Prior to seeding the gel, iCFs were cultured for 48 h with DOX-conditioned media or a blank control, and iCMs were cultured under normal conditions. After 3D bioprinting, constructs were allowed to adjust in a 1:1 mixture of CM(+) and DMEM complete for 14 days. Constructs were then separated into 3 groups: control iCFs treated with regular media (control), control iCFs treated with media supplemented with 10 nM DOX (DOX), and DOX pre-treated iCFs treated with regular media (pre-treated).

Beating behavior was assessed after 48 h media treatment. After 48 h, a significant decrease in beating velocity was observed in both the DOX (22%) and pre-treated (43%) groups, with the pre-treated group reducing beating velocity substantially, but not significantly, more (Figure 1F, left). Additionally, while the DOX treated group showed significantly increased cell death, the pre-treated group did not (Figure 1F, right). In addition to significant disruptions to beating velocity and rate, DOX pre-treatment also substantially disrupted the beating regularity of the 3D models (Figure 1G), similarly to what was observed in 2D culture, while the control group showed even greater regularity in beating compared to 2D culture. Subsequent FFT and sine wave decomposition further support this interpretation (Supplemental Figure S2).

### DOX-EVs Alone Induce Metabolic Dysfunction Expected in DOX-Cardiotoxicity

Following regular cell culture assay, the mitochondrial respiration and associated functions of cells were analyzed via Seahorse. All three cell types: iCMs, iCFs, and iECs, were seeded on Seahorse-compatible 96-well plates and allowed to settle. After attachment, cells were treated for 48 h with DOX media, a PBS blank, or DOX EV media where the EVs originated from either DOX-treated CFs (CF DOX-EVs) or DOX-treated MFs (MF DOX-EVs). This was done to see if DOX-EVs from non-cardiac tissues could induce similar dysfunction to those from CFs. Five major categories were considered: basal respiration (showed no significant difference in any group), maximal respiration (MR, Figure 2A), ATP-production coupled respiration (ACR, Figure 2B), proton leak (PL, Figure 2C), and spare respiratory capacity (SRC, Figure 2D). In iCMs and iCFs, both the MR and SRC were significantly decreased by all treatment groups, demonstrating that, independent of DOX treatment, DOX-EVs are sufficient to induce significant metabolic distress in cardiac cells, and that the cells contributing to these effects do not necessarily need to be of cardiac origin. This is in conjunction with a significant increase in ACR in iCFs subjected to both DOX and CF DOX-EV treatment, and in iCMs subjected to CF DOX-EV treatment but not DOX treatment alone. The combination of decreased MR and SRC (Figure 2A, D) with increased ACR (Figure 2B) is characteristic of DOX-associated mitochondrial dysfunction in cardiomyocytes (CMs)^22,23^ and with larger-scale anthracycline-induced progressive cardiotoxicity^24^. This is consistent with the results of the 2D and 3D models. MF DOX-EVs, alternatively, significantly decreased PL, a measure of mitochondrial inefficiency (Figure 2C), which may indicate a different, though still disruptive, influence from non-cardiac tissues. On the other hand, iECs experienced significantly increased MF and SRC under all conditions. This combined with the increase in ACR, suggests either increased overall mitochondrial activity or an increase in the number of mitochondria. With the increase in PL as well, however, the mitochondrial efficiency also likely decreased significantly. This pattern aligns with the 2D model, which showed a non-significant increase in ROS alongside a significant reduction in overall mitochondrial activity (Figure 1E)

**Figure 2:**
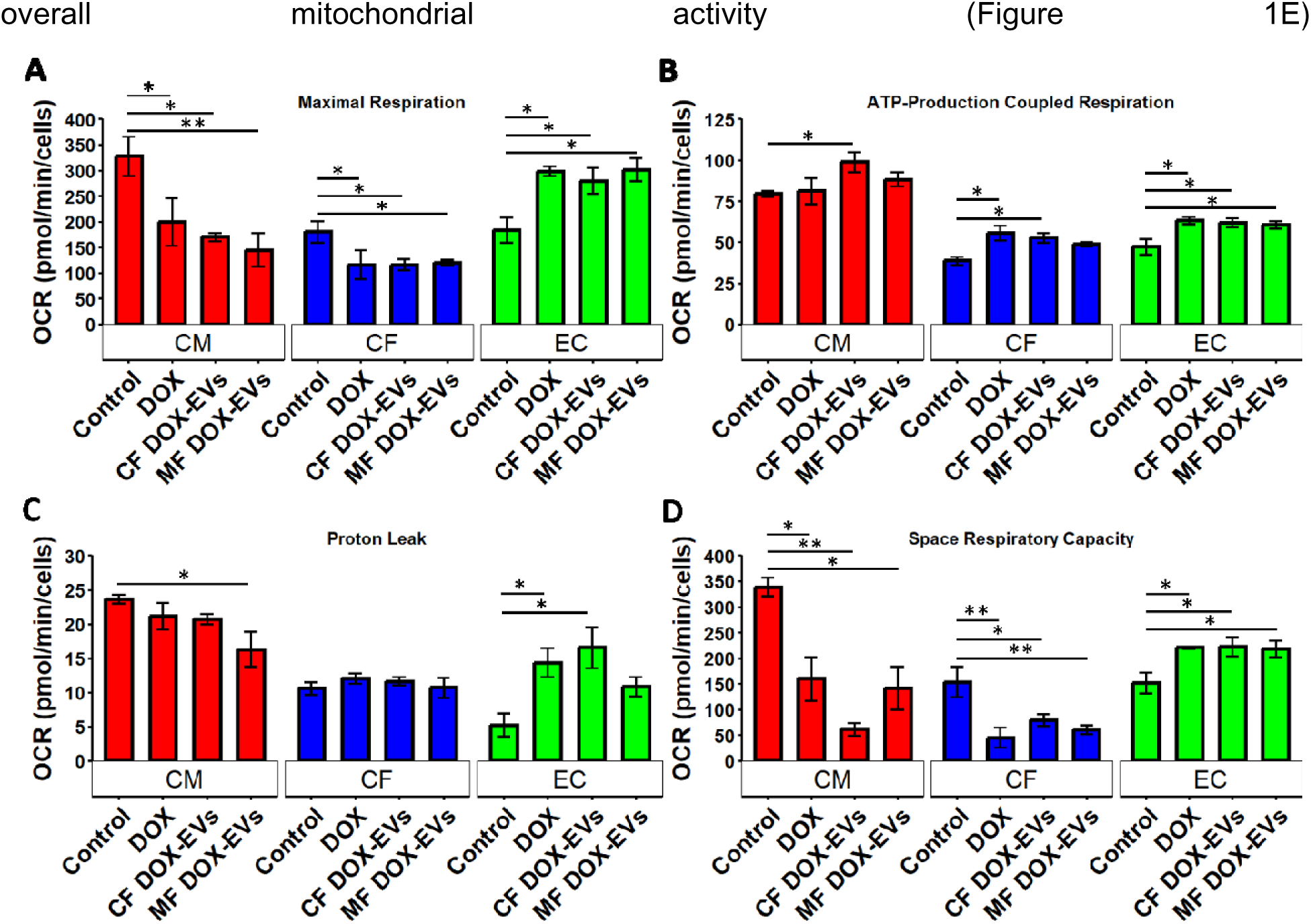
DOX-EVs Induce Metabolic Dysfunction Similar to Direct DOX Treatment. Seahorse MitoStress assay results for (A) maximal respiration, (B) ATP-production coupled respiration, (C) proton leak, and (D) spare respiratory capacity after 48 h of treatment. n ≥ 3 for all groups tested, * p < 0.05, ** p < 0.01, assessed by one-way ANOVA with Tukey’s post-hoc

### DOX Treatment Induces Pathology-Associated Shift in EV Characteristics

To establish that DOX-treatment alters the paracrine signaling behaviors of cardiac cells (Figure 3A), EVs were collected from conditioned, exosome-free media from iCMs, iCFs, and iECs treated with a blank control or 10 nM DOX. To demonstrate that changes in EV populations were not a result of direct DOX export in EVs, we first performed UV-Vis spectrophotometry to assess the presence of DOX in a PBS blank, DOX-conditioned media, and EVs from both control and DOX-treated iCMs and iCFs (Figure 3B, Supplemental Figure S3). We observed more than 2-order of magnitude difference between DOX-conditioned media and any other group, and that all EV groups were nearly identical. Furthermore, the absorbance spectra for all EVs were virtually indistinguishable, whereas the spectra from DOX-conditioned media was clearly distinguishable with peaks not present in any other spectra (Figure 3B).

**Figure 3:**
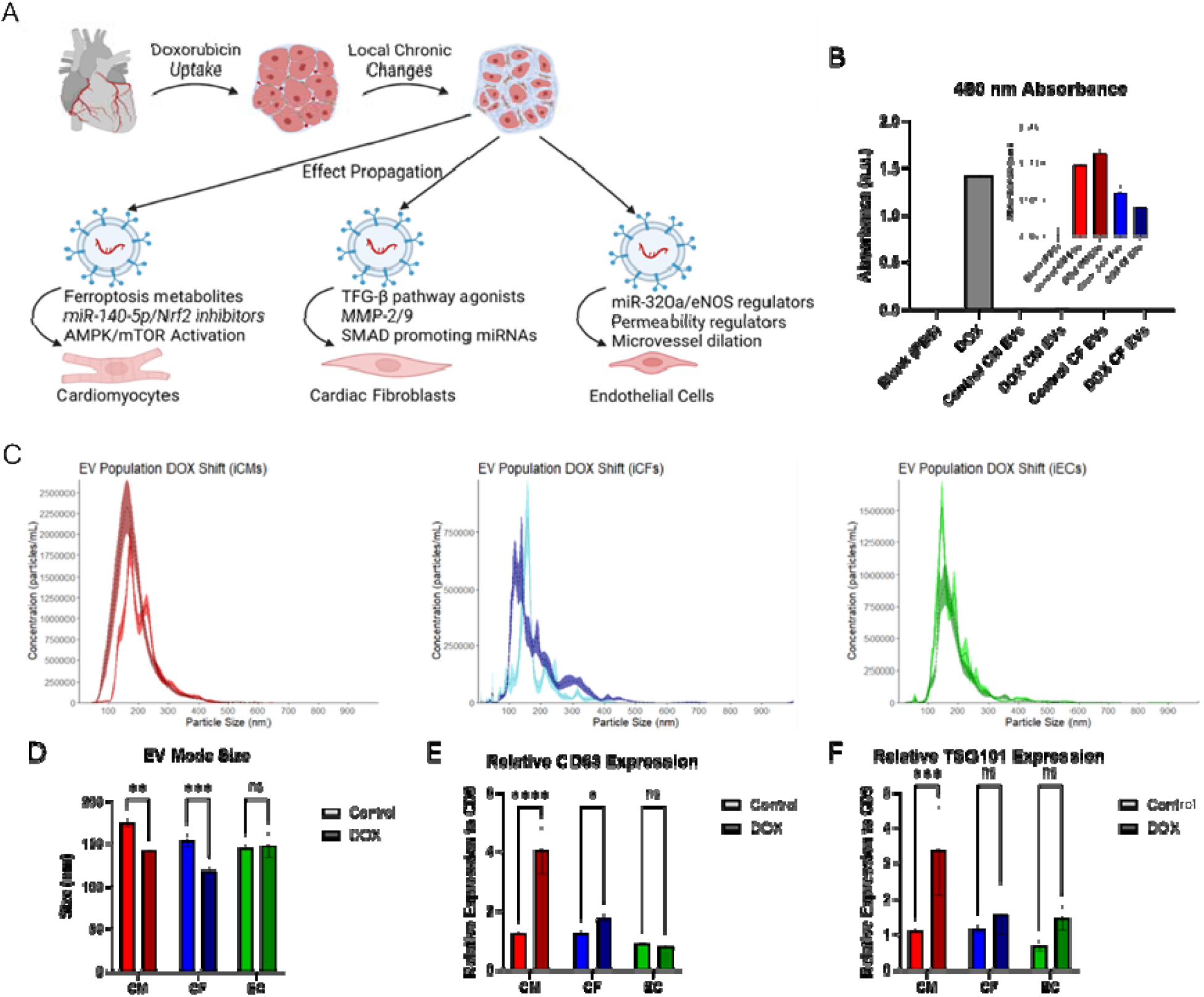
DOX-EVs show disease-like morphology and protein profiles without acting as direct doxorubicin carriers. (A) Diagram briefly showing the process of events by which DOX cardiotoxicity may be propagated by local paracrine signaling. (B) The absorbance of a PBS blank, DOX-conditioned media, and EVs from control and DOX-treated iCMs and iCFs quantified by UV-Vis spectrophotometry for 480 nm. (C) Nanoparticle tracking analysis (NTA) with error area for EVs from control or DOX-treated iCMs (red, left), iCFs (blue, middle), or iECs (green, right), with (D) subsequent quantification of mode shift in each population. Quantification of the western blot band intensity of CD63 (E) or TSG101 (F) relative to CD9 intensity for EVs obtained from control or DOX-treated iCMs, iCFs, and iECs. Data are presented as the mean ± standard deviation. n ≥ 3 for all groups tested, * p < 0.05, ** p < 0.01, *** p < 0.005, **** p < 0.001 assessed by one-way ANOVA with Tukey’s post-hoc for (D), (E), and (F).

Following this, the size profile of isolated EVs was assessed via Nanoparticle Tracking Analysis (NTA) (Figure 3C). This showed that EVs from DOX-treated cells tended be smaller than those from control cells in iCMs (Control, mode: 174 nm; DOX, mode: 141 nm; p < 0.01) and iCFs (Control, mode: 153 nm; DOX, mode: 118 nm; p < 0.005), but not in iECs (Control, mode: 144 nm; DOX, mode: 147 nm) (Figure 3D). All EVs measured fall within the expected range for EVs (< 200 µm diameter), and all populations have calculated PDI below 0.2, indicating that the EV populations are mostly monodisperse and therefore not splitting into easily separable sup-populations.

Western blot was performed to identify characteristic exosome markers CD9, CD63, and TSG101 (Supplemental Figure S4). This was done to both identify whether the population contained exosomes, as well as to assess the relative expression of CD63 and TSG101 to CD9 (Figure 3E-F), which we have previously suggested may be correlated with damage or other dysfunction in the myocardium^14^. These results showed that the isolated EV populations did contain exosomes and showed significant changes in some surface marker expression. Compared to the blank control, relative CD63 expression was increased in DOX-treated iCMs (p < 0.001) and iCFs (p < 0.05), and relative TSG101 expression was increased in DOX-treated iCMs alone (p < 0.005), in line with our previous findings regarding pathology-related changes in EV tetraspanin expression^25^.

### DOX Treatment Substantially Alters EV miRNA Profile of Cardiac Cells

miRNA profiling via Nanostring analysis revealed highly upregulated clusters of exosomal miRNA populations in EVs from DOX-treated cells relative to control cells (Figure 4A). Interestingly, despite this separation between control and DOX-treated cell EVs, the miRNA populations of EVs from control iCMs and DOX-treated iCFs clustered together, though the miRNA profiles are still notably distinct. From over 800 miRNAs profiled, unsupervised analysis revealed 146 miRNAs (Supplemental Table S1) that were selected as being meaningfully altered as a result of DOX treatment (Figure 4B). These miRNAs clearly clustered between control and DOX treated cell EVs, and more miRNAs were upregulated in the DOX treated cell EVs than in those from control cells. There was also substantial overlap between both DOX treated cell EV profiles and the control CM profile, though many of these miRNAs mapped to normal cardiac processes.

**Figure 4:**
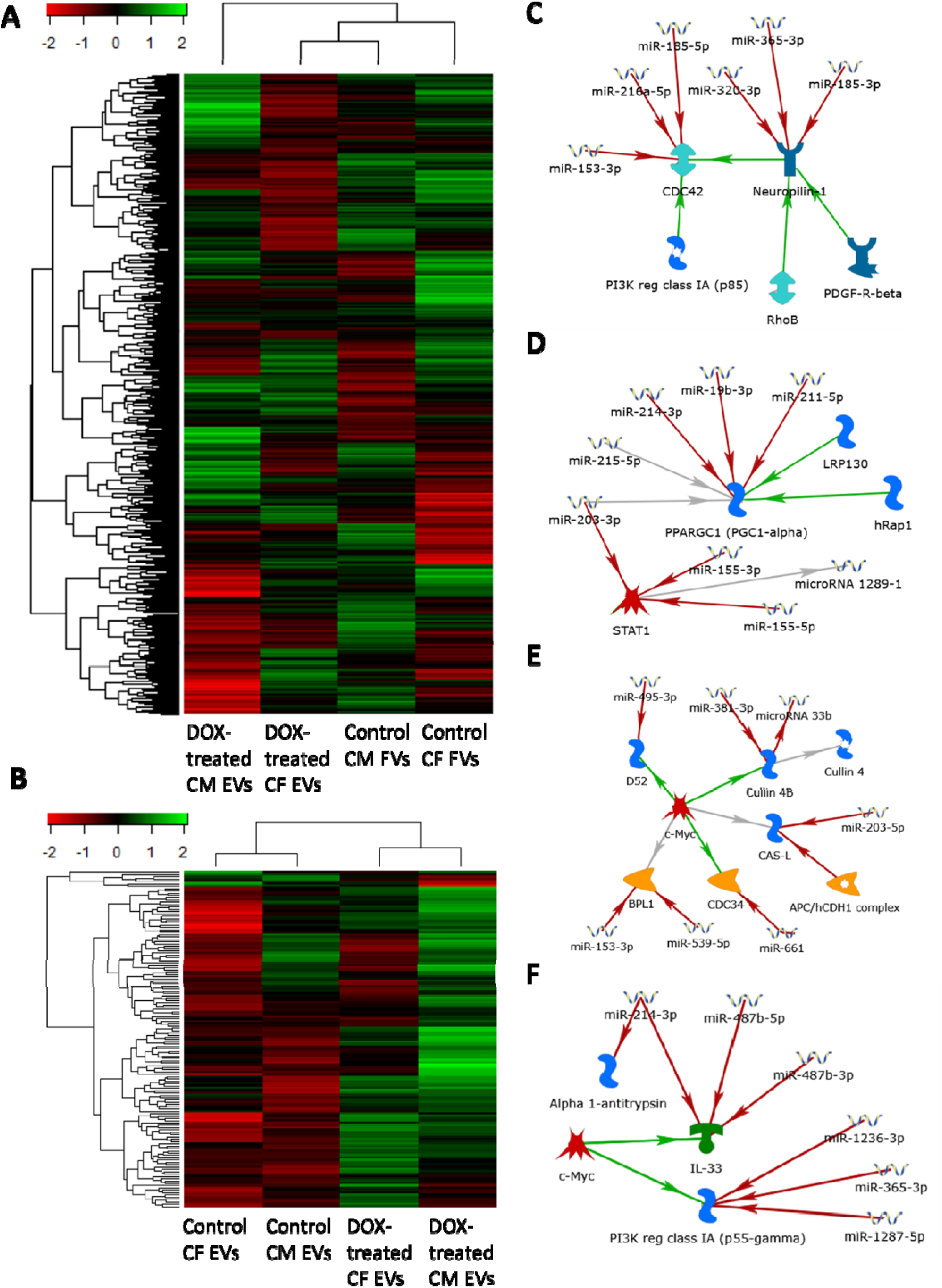
DOX-EVs demonstrate a differential miRNA profile and target disease pathway regulation. (A) Heatmap showing the full miRNA profiling of isolated EVs from iCMs and iCFs after 48 h treatment with DOX-conditioned media or a blank control. (B) miRNA profiling results for 145 identified targets for downstream analysis. (C-F) Results of MetaCore pathway analysis for the identified target miRNAs. Identified pathways were considered for analysis only for p-value < 0.05.

### miRNAs upregulated in DOX-cell EVs are associated with cardiotoxicity pathways

To assess potential overlap and pathways of interest, MetaCore pathway analysis software was used to build networks for the 145 identified distinctive miRNAs. This network analysis identified several pathways of interest that were regulated by two or more of the identified target miRNAs, often with multiple miRNAs regulating a single target or series of targets. The most involved pathway identified was that of CDC42 and upstream Neuropilin-1 (Figure 4C, full pathway: Supplemental Figure S5A), a cell proliferation pathway often desirably hindered by chemotherapy and is vital in some cancer progression^26^, and utilized 6 miRNAs (Supplemental Table S2). This indicates that miRNAs delivered by sEVs from DOX treated cells continue to promote anti-cancer effects. Also notable was the regulation of PPARGC1-a (Figure 4D), the “master regulator” of mitochondrial biogenesis^27^. PPARGC1-a is also responsible for translation of mechanical stimuli to mitochondrial biogenesis^28^ and the promotion of M2-phenotype macrophage polarization^29^ in the heart, and downregulation of this gene as a result of genetic mutation was found to increase risk of left ventricular diastolic dysfunction^30^. This interaction was regulated by 5 miRNAs (Supplemental Table S2). Both of these interactions were found in the same pathway (Supplemental Figure S5A). Another pathway of interest revealed four interactions downstream of c-Myc (Figure 4E, full pathway: Supplemental Figure S5B). Of interest in these interactions were the upstream and downstream regulation of Cullin 4B, a known effector of cardiac antioxidant pathways and sarcomere quality control and which the downregulation of is associated with heart failure^31^. This interaction involves 2 miRNAs (Supplemental Table S2). The other interaction of major interest is the inhibition of CAS-L by miR-203-5p. CAS-L is a cardiac redox agent via MICAL1 involved in the sensing and binding of Ca^2+^, the inhibition of which is associated with ventricular tachycardia^32^. Finally, two more interactions downstream of c-Myc in a different pathway were identified (Figure 4F). The first interaction is the downregulation of a PI3K component by 3 miRNAs (Supplemental Table S2). Although PI3Ks are desirable targets for arresting breast cancer, PI3Ks are also essential cardioprotective agents and the inhibition of them by pharmaceuticals is frequently met with off-target cardiotoxicity and arrhythmia^33^. PI3Ks are an expected, though not well mapped-out, target of anthracyclines and the application of DOX with PI3K supplementation through pharmaceuticals or ischemic preconditioning has alleviated some of the cardiotoxic effects of DOX treatment in animal models^34^. The other interaction of interest was the inhibition of IL33 by 3 miRNAs (Supplemental Figure S5C) and subsequent inhibition of Alpha 1 antitrypsin (AAT). IL33 has recently been implicated an essential component of a mechanically sensitive CF cardioprotective paracrine signaling machinery, protecting against hypertrophy, fibrosis, and heart failure^35^ and AAT deficiency is associated with many CVDs and systemic failure of the heart through unknown mechanisms^36^.

### Distinct Plasma EV miRNA Profiles Differentiate High- and Low-Risk Cardiotoxicity Patients

EVs were isolated from blinded clinical plasma samples of patient cohorts classified as high or low risk for DOX-induced cardiotoxicity to profile their miRNA content and identify molecular signatures associated with cardiotoxicity risk. The patient samples clustered distinctly by risk category, revealing that high-risk patients exhibit plasma EV miRNA signatures similar to those observed in DOX-treated cell culture models (Figure 5A). To pinpoint the miRNAs most responsible for this separation, we focused on the top 10% of miRNAs (n = 80) showing the highest percent coefficient of variance (%CV; red dashed line, Figure 5B). Among these, 23 miRNAs displayed particularly pronounced variability across five distinct cohorts (Supplemental Table S3), as visualized in the heatmap and clustering analyses (Figure 5A). These cohorts consisted of the identified 23 miRNA targets and adjacent miRNAs (mostly consisting of others of the 80 most enriched miRNAs), and were distinguished to the right of the heatmap as follows: cohort 1 – grey; cohort 2 – orange; cohort 3 – blue; cohort 4 – green; cohort 5 – purple (Figure 5A, Supplemental Table S3). Importantly, 35 of the miRNAs enriched in patient EVs closely overlapped with miRNAs previously identified in DOX-treated cell culture EVs (Figure 5C), and were largely enriched in patients with high risk of cardiotoxicity (Figure 5D).

**Figure 5:**
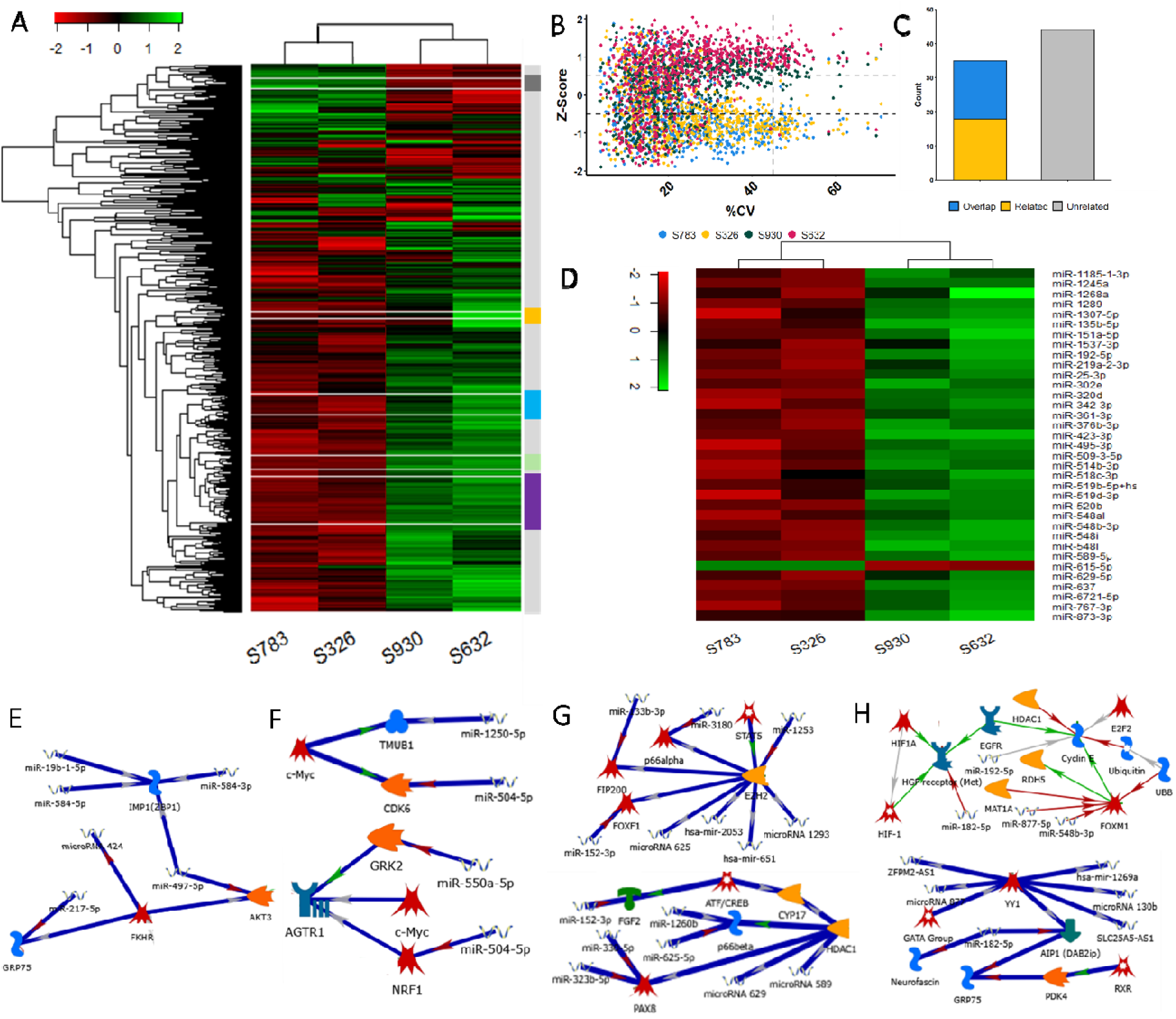
Clinical samples mimic DOX-miRNA enrichment in low vs high DCT-risk patients. (A) Heatmap showing hierarchical clustering of the top 10% most distinguishing miRNAs (n = 80 by % coefficient of variance (%CV) across four blinded patient samples (S783, S326, S930, S632). Five major expression clusters are color-coded on the right. (B) Scatter plot of miRNA z-scores versus %CV, with the top 10% most distinguishing miRNAs outlined (blue box, right of red dashed line). Each dot is color-coded by patient sample. (C) Bar graph quantifying the overlap between high-%CV plasma EV miRNAs and those identified in DOX-treated tissue-engineered models: fully overlapping (blue), partially related (orange), or unrelated (gray) with a heatmap representing relative enrichment of these miRNAs in each patient sample (D). MetaCore pathway analysis was performed to identify relevant pathways for identified miRNA targets of each cohort, and pathways relating to chemotherapy-induced cardiotoxicity were identified in cohort 1 (E), cohort 2 (F), cohort 3 (G), and cohort 4 (H).

### Enriched miRNAs in High DCT Risk Patients Regulate Cardiotoxic Pathways

Pathway enrichment analyses indicated that many of these miRNAs (both shared and unique) are implicated in pathways governing cardiac homeostasis, chronic CVD development, and the progression of cardiotoxicity (Figure 5E-H), though analysis of cohort 5 showed little specificity for cardiotoxicity and was more closely related to general dysregulation of cardiac homeostasis (Supplemental Figure S6), possibly indicating an overselection of targets. Specifically, miRNAs from cohort 1 miR-584-3p^37,38^and miR-217-5p^41,42^ have the most direct reported links to DCT, whereas miR-497-5p^43,44^ and miR-1262^45,46^ have been independently associated with both CVD-related pathways and anti-tumor responses (Figure 5E, Supplemental Figure S6A). MetaCore analysis also identified significant co-operative regulation of DCT-related pathways, including GRP75^47^ and AKT^48,49,50,51,52^, known to be recruited during DOX-induced stress signaling, and downstream FOXO^53,54,55^, linked to metabolic disruption and apoptosis after DOX-mediated disruption of AKT, in parallel with disruption of ZBP1/IMP1^56,57^, a disruption which has been shown to be necessary for DCT onset in animal models (Figure 5E). Additionally, cohort 1 miRNAs also disrupted c-Jun related signaling, shown to cause metabolic stress and pathological remodeling in the heart, via AP-1^58,59^, c-Abl^60^, and C/EBP^61^, with c-Abl being directly implicated in DCT onset (full pathway: Supplemental Figure S6A).

The miRNAs clustered in cohort 2 were implicated in chronic CVD and fibrotic remodeling and are being investigated as targets for intervention, including miR-504^78,79^ miR-553^76,77^, miR-604^81^, miR-595^82^, and miR-545-3p^83^ and miR-1250-5p^80^ (Figure 5F, full pathways: Supplemental Figure 6B-C). While the cluster 2 miRNAs were not as directly implicated in the onset of DCT in existing literature, MetaCore analysis revealed several major DCT-related proteins being co-operatively regulated by DOX-shifted miRNAs across two overarching pathways, both downstream of or in parallel to c-Myc. Both CDK6^62,63^ and TMUB1^64,65,66^ have been linked to chemotherapy-related cardiotoxicity, specifically via p53 regulation and metabolic stress, and in this pathway are in parallel downstream of c-Myc (Figure 5F, upper). Additionally, in parallel with the above pathway, NRF1^67,68^, GRK2^69,70,71^, and AGTR1^72^ signaling were also disrupted, the disruption of each of which has been directly linked to DOX-induced damage, particularly in suppression of mitochondrial function and p53 regulation (Figure 5F, lower, full pathways: Supplemental Fig. 6B-C).

The miRNAs clustered in cohort 3 were largely linked to DOX-induced oxidative stress and subsequent CM death, with miR-152-3p^73,74,75^ being a well-known individual miRNA target in DCT studies. MetaCore analysis of cluster 3 linked these miRNAs to largely fibrosis and chronic CVD-related pathways. In particular, the disruption of EZH2^84,85^, p66a^86,87,88^, FOXF1^89,90,91^, and FIP200^92,93^ have been mechanistically linked to the onset of excessive fibrosis in a number of CVDs via both pathway regulation and gene activation, with p66a being actively explored as an anti-fibrotic drug target (Figure 5G, upper). In parallel, however, cluster 3 was also highly involved in pathways directly related to chemotherapy-induced cardiac fibrosis, with both HDAC1^94,95^ and p66b^96,97^ being implicated (though for trastuzumab, not DOX), and both PAX8^98,99^ and ATF/CREB/FGF2^100^ axis disruption exacerbating any induced fibrosis (Figure 5G, lower, full pathways: Supplemental Fig. 7A-B).

The miRNAs clustered in cohort 4 were also largely implicated in the onset of DCT, such as miR-182-5p^101^, via regulation of major DCT nodes. MetaCore analyses linked this cluster to highly interconnect nodes associated with a wide array of chemotherapy induced cardiac toxicity and fibrosis, including UBB^102,103,104^ FOXM1^105,106,107,108^ Cyclin E kinase^109,110^ HGFR/HIF1^111,112,113,114^, all of which are major targets of investigation for the onset of chemo-CT, and, interestingly, all of which have been shown to be induced to pathology via irregular miRNA signaling (Figure 5H, upper). More directly, YY1^115,116^ AIP1^117^, and GRP75^118^ have been directly implicated in DCT, poor ion handling, and DCT-related heart failure, with AIP1 being known to be directly regulated via EV-chauffeured miRNAs to manipulate pathway activation (Figure 5H, lower). While further biological replicates and direct, targeted clinical studies are required to validate both these markers and the observed trends and interactions, these preliminary data suggest that plasma EV miRNA profiling could serve as a promising biomarker tool for identifying patients at risk for DOX-induced cardiotoxicity, with translational alignment between *in vitro* and clinical settings.

## DISCUSSION

In this study, we evaluated the effects of DOX treatment and treatment with EVs derived from DOX-treated cells side-by-side in both 2D and 3D *in vitro* models. These models primarily consisted of iCMs and iCFs, cells which make up a vast majority of the heart^20^, and iECs, which are commonly used in stem cell-derived cardiac tissue models to emulate vasculature^25^. In all cell types we evaluated ROS prevalence and mitochondrial activity, and for cardiac cells specifically we evaluated metrics of cardiac health including beating velocity in iCMs and pro-fibrotic transdifferentiation in iCFs. The health and survivability of iCMs was then evaluated again in a 3D bioprinted cardiac tissue model containing iCFs that had been pre-treated with either DOX or a PBS blank, and iCMs and allowed to grow for 14 days. These constructs were then treated with either control media (for both the control and pre-treated groups), or DOX-containing media for 48 h, after which the CM beating, beating regularity, and viability of the cells was analyzed. Following this, the effects of direct DOX treatment and DOX-EVs from cardiac and breast fibroblasts on mitochondrial respiration in 2D cultures were assessed using the Seahorse MitoStress assay, which quantifies oxygen consumption rates (OCR) to derive metrics of mitochondrial function. After establishing the effects of DOX-EV treatment relative to DOX, we evaluated the size, morphology, and surface protein profiles of EVs derived from iCMs, iCFs, and iECs with or without DOX treatment.Once complete, the miRNA cargo of the EVs and DOX-EVs from iCMs and iCFs were profiled using Nanostring miRNA profiling and investigated for downstream interactions pertaining to DOX cardiotoxicity via MetaCore pathway analysis. Following this, we isolated EVs and their miRNA cargo from the plasma of patients receiving DOX treatment and assessed for high or low risk of cardiotoxicity, and performed Nanostring miRNA profiling for each subject. We were able to successfully bifurcate these populations based off of the miRNA profiles obtained and compare distinguishing miRNAs from the clinical samples with those from our models. From this, we were able to identify 5 major clusters of miRNAs highly related to the onset of DCT and further evaluate the activities of these miRNA clusters via MetaCore pathway analysis. From these data, we identified miRNAs which could be obtained from plasma which may serve as a basis to develop quantitative assays for the early detection of DCT in chemotherapy patients, and identified major miRNA regulated pathways which may bolster the development of future intervention strategies to inhibit or prevent the onset of DCT.

Herein, we have shown, for the first time in literature, that EVs secreted from cardiac stromal cells, in this case iCFs, can independently replicate the effects of DOX cardiotoxicity without ever directly exposing the target cells to DOX. While the assays performed in this study are by no means exhaustive, in iCMs it was possible to replicate increased oxidative stress, whether that be through increased ROS or decreased antioxidants, and significantly dysfunctional beating behaviors reminiscent of DOX cardiotoxicity but without directly using DOX. These results were consistent in iCFs as well, which showed distinctly pro-fibrotic and oxidatively stressed behaviors which were not the result of excessive cell death. Furthermore, these effects did not require deliberate EV exposure via conditioned media, which could introduce trace amounts of DOX if EVs are not isolated or washed appropriately. In the 3D culture, the iCFs which were exposed to DOX were washed thoroughly before seeding and allowed 14 days to settle, which, evaluating the substantial increase in cell death when iCFs were exposed to DOX, would be sufficient time for any residual DOX to induce significant amounts of cell death. However, no significant change in cell death was observed in the pre-treated models and the iCMs within those models exhibited significant dysfunction without ever having been directly exposed to DOX. This strongly suggests that DOX cardiotoxicity can be induced over time via affected CFs in the heart, rather than directly from DOX itself. This may help explain a central paradox of anthracycline cardiotoxicity, namely, how the heart can be so strongly affected given the known rapid clearance time of DOX (terminal half-life of 30 h and systemic clearance within 48 h).

Additionally, in iCMs and iCFs, but not iECs, CF DOX-EVs were able to induce similar metabolic dysfunction to direct DOX treatment, and this dysfunction was consistent with what is observed in DOX cardiotoxicity in *in vivo* models^22^, though this effect was not consistent with MF DOX-EVs. This discrepancy may be due to mammary fibroblasts having a slightly different, less cardiotoxic response to DOX treatment than cardiac fibroblasts, or it may be due to differences between primary and iPSC-derived fibroblast behaviors and paracrine signaling. However, it remains interesting that CF DOX-EVs were able to recapitulate the impact of DOX treatment on both iCMs and iCFs. These data, taken together with the 2D and 3D model data, suggest that DOX cardiotoxicity can, to a degree, become self-propagating by inducing DOX-like dysfunction in nearby CFs. Thus, even a brief initial exposure of the heart to DOX may be sufficient to affect a subset of cells and substantially elevate the long-term risk of cardiotoxicity.

The effects of DOX treatment on iCM and iCF paracrine signaling further implicate EVs as key contributors to DOX-induced cardiotoxicity. As EVs are essential agents for maintaining tissue homeostasis in the heart^119^, significant alterations in their size or cargo can have a profound impact on the development of chronic CVDs^120^. Furthermore, a significant decrease in EV size has previously been linked to chronic CVD^120^, as has alterations in the surface tetraspanin web^25^. Moreover, these changes were not a result of EVs being repurposed for DOX export from cells, although EVs are sometimes deliberately loaded with DOX to generate an endogenous vehicle for drug delivery^121^. These data, however, support preliminary conclusions from the cell culture and Seahorse MitoStress assay data. As DOX-EVs do not contain DOX itself, there must be some other alteration occurring in or to the produced EVs which are inducing the observed DOX cardiotoxicity-like changes.

Full miRNA profiling of EVs from control iCMs and iCFs compared to EVs from iCMs and iCFs treated with DOX for 48 h provides interesting insight into what specifically these changes may be. While the full profile does not provide much useful information, unsupervised analysis of the dataset revealed 145 potentially relevant miRNA targets, and, when clustered, these 145 targets clearly delineate between control and DOX-treated cell EVs. More interesting, perhaps, is the downstream activities of the identified EVs which were assessed by MetaCore. The pathways affected, when in the context of cancer, are useful pathways to utilize to dysregulate redox balance, immunomodulation, and disruption of regular mitochondrial functioning, though the mechanisms are not well understood, and in fact partially converged (with 6 miRNAs “cooperating” on this pathway) on CDC42 and upstream Neuropilin-1, a cell proliferation pathway often desirably hindered by chemotherapy and is vital in some cancer progression^26^. When these same pathways are dysregulated in the heart, however, can cause significant damage to the surrounding tissue and microenvironment by way of oxidative stress^31,33^, disruption of ion handling and beating regulation^30,32^, maladaptive remodeling and pathology-associated changes in paracrine signaling^35^, and other heart failure-associated effects.

Interestingly, this connection may answer more about how DOX cardiotoxicity works than just how the condition manifests long-term. While it is well established that DOX induces dysfunction in CMs by damaging the mitochondria and often causing mitochondrial depolarization or ferroptosis, the precise mechanisms by which this occurs in CMs are essentially unknown^122^. Furthermore, the general understanding of the role that mitochondria play in maintaining cardiac health has recently grown by leaps and bounds. In particular, mitochondrial imbalance has been a recent area of interest, especially for cancer and chemotherapy-related CVDs^123^. Mitochondrial imbalance is when CMs, which have a relative overabundance of mitochondria compared to most other cells in the body, are incapable of disposing of dysfunctional mitochondria either due to dysfunctional mitochondrial disposal or increased mitochondrial damage. One way that this can manifest is observed through decreased mitochondrial capacity, but increased mitochondrial activity, as was observed in the cell culture and metabolism assays with CMs treated with DOX and DOX-EVs. Furthermore, it has recently been suggested that DOX-induced cardiomyopathy stems largely from disrupted redox circuits in cardiac cells, altered metabolic activity and resulting mitochondrial stress, and dysfunctional ion handling, through secondary interactions that result from DOX exposure rather than direct DOX interactions^124^, though the precise mechanisms by which these may occur is unknown. It is, however, compelling that the top common results for pathways which the identified target miRNAs are involved in directly correlate to those which are now hypothesized to drive DOX-induced cardiotoxicity.

In conclusion, we have established, for the first time in literature, that DOX-induced cardiotoxicity can be induced in cardiac cells in the absence of DOX by using EVs from cells which have previously been exposed. These effects are consistent in 2D and 3D cell culture models, and mimic the expected dysfunction of DOX cardiotoxicity. Furthermore, we have demonstrated that DOX treatment significantly alters the paracrine profile of cardiac cells, both in terms of size and miRNA cargo, and that these altered miRNA cargos are highly and jointly involved in processes which are hypothesized to be major drivers of DOX-induced cardiotoxicity. To build upon these findings, we subsequently profiled plasma EVs from breast cancer patients at high or low risk for DOX-induced cardiotoxicity and observed similar miRNA alterations, further supporting the translational relevance of our in vitro models and the potential of EV miRNAs as predictive biomarkers. Our analysis of plasma EV miRNAs from patient cohorts further underscores the translational potential of EV cargo profiling as a biomarker strategy for DOX-induced cardiotoxicity. The distinct clustering of high- and low-risk patients, coupled with the substantial overlap between patient-derived EV miRNAs and those identified in DOX-treated *in vitro* models, provides compelling evidence that EV miRNA signatures reflect both the initiation and propagation of cardiotoxic signaling. Importantly, several of the enriched miRNAs were linked to pathways central to cardiac homeostasis, mitochondrial regulation, and fibrotic remodeling, suggesting that these circulating EVs may not only serve as indicators of cardiotoxicity risk but also provide mechanistic insight into the pathophysiology of chronic DOX injury.

While this study provides an excellent first step into understanding the interplay between local paracrine signaling and both acute and chronic DOX cardiotoxicity, this study has several limitations. First, the endothelial cells utilized in this study are not cardiac specific, and as such may exhibit different behaviors than cardiac specific endothelial cells may. Second, the cell assays performed in this study are fairly preliminary, although they do cover a wide breadth of cells. To more completely compare and contrast the impact of DOX and DOX EVs on cardiac cells, future studies would be prudent to focus more completely on a single cell type or otherwise a more biomimetic model, such as microtissues or heart-on-chip devices, to provide a more complete picture of what interactions may be occurring. Lastly, this study did not completely investigate the interactions between non-cardiac tissues exposed to DOX. While mammary fibroblasts were briefly investigated and showed notably different interactions, it would be beneficial in future studies to evaluate the impact of DOX EVs from other tissues on cardiac health. Great efforts are being made to target anthracyclines to the desired tissues in order to circumvent cardiotoxicity, and understanding how these tissues may then downstream interact with the myocardium would be extraordinarily useful. In addition, while the clinical EV profiling offers strong translational support for our findings, the modest sample size limits broader generalization and highlights the need for validation in larger patient cohorts. Moreover, future studies incorporating intermediate models, such as animal models or ex vivo platforms, could help capture systemic factors and strengthen the mechanistic link between EV signaling and cardiotoxicity observed in both engineered tissues and clinical samples. Despite these limitations, this study has been an incredibly successful first step down a long road of understanding the interactions between EVs and DOX cardiotoxicity, and this and future studies will help finally elucidate the mechanisms by which DOX-induced cardiotoxicity occurs.

## MATERIALS & METHODS

### Culture of Human Induced Pluripotent Stem Cells (hiPSCs)

DiPS 1016 SevA hiPSCs, which were derived from human skin fibroblasts, were cultured on Geltrex (1% Invitrogen, USA)-coated culture flasks in mTeSR (StemCell Technologies, Canada) media supplemented with 1% penicillin (VWR, USA) under standard culture conditions from passages 40-50.

Cells were passaged at 80% confluency. To passage the hiPSCs, cells were detached using Accutase (StemCell Technologies, Canada) and seeded onto Geltrex-coated cell culture well plates or split between cell culture flasks. Seeding was performed in mTeSR media supplemented with Rho-associated, coiled-coil containing protein kinase (ROCK) inhibitor (5 μM, StemCell Technologies, Canada). For differentiation, cells were cultured until 95% confluency before starting a protocol.

### Differentiation and Culture of hiPSC-derived Cardiomyocytes (iCMs)

Differentiation of hiPSCs to iCMs was adapted from a previously established protocol. Briefly, when hiPSCs were ready for differentiation the media was changed to RPMI Medium 1640 (Life Technologies, USA) supplemented with B27 without insulin (2%, Invitrogen, USA), and beta-mercaptoethanol (final concentration of 0.1 mM, Promega, USA) (CM (-)) with the addition of Wnt activator, CHIR99021 (CHIR) (10 μM, Stemgent, USA) (Day 1). Exactly 24 h later, media replaced with CM (-) with CHIR (2 µM) (Day 2), and again another 24 h later (Day 3). Exactly 24 h later, media was changed to CM (-) with Wnt inhibitor IWP-4 (5 μM, MA, USA) (Day 4). Exactly 48 h later, media was changed to CM (-) (Day 6). Exactly 72 h later, media was changed to RPMI Medium 1640 supplemented with B27 (2%, Invitrogen, USA), and beta-mercaptoethanol (final concentration of 0.1 mM) (CM (+)) (Day 9). Following this, every 3 days media was changed using CM (+). Cultures typically began to beat by day 21 of this protocol, as reported previously^125^, and continued to be cultured in CM (+) until use.

### Differentiation and Culture hiPSC-derived Cardiac Fibroblasts (iCFs)

Differentiation of hiPSCs to iCFs was adapted from a previous protocol^126^. Briefly, when hiPSCs were ready for differentiation the media was changed to CM (-) with 10 µM CHIR (Day 1). After 24 h, media was changed to CM (-) without CHIR (Day 2). After 24 h, media was replaced with CFBM media supplemented with fetal bovine serum (FBS) (Gibco) and 75 ng/mL fibroblast growth factor (FGF) (Day 3). Media was refreshed every 48 h until day 20. After day 20, differentiated iCFs were detached from the cell plate with trypsin-EDTA (0.25%, Stem Cell Technologies, Canada) and seeded into fibronectin (Sigma Aldrich, USA)-coated cell culture flasks. Seeded iCFs were from then on cultured in Dulbecco’s Modified Eagle Medium (DMEM) (Thermo Fisher) supplemented with 10% FBS, 1% penicillin/streptomycin (P/S) (Life Technologies), henceforth called DMEM Complete, and 3 µM SD208, a TGF-b receptor I kinase inhibitor (Sigma Aldrich) on fibronectin-coated cell culture flasks. Cells were cultured with SD208 supplement to inhibit transdifferentiation, and then used between passage 4 and 10 without SD208. Human mammary fibroblasts (hMFs) were received from a collaborator and cultured using the above protocol.

### Differentiation and Culture hiPSC-derived Endothelial Cells (iECs)

Differentiation of hiPSCs to iECs was adapted from a previous protocol^125^. Briefly, when hiPSCs were ready for differentiation the media was changed to a 1:1 mixture of DMEM to F12 with Glutamax and Neurobasal media supplemented with N2 (1%, Life Technologies, USA), B27 (2%), CHIR (8 μM) and bone morphogenic protein 4 (25 ng/ml, R&D Systems, USA) (Day 1). After 72 h, media was replaced with StemPro-34 SFM medium (Life Technologies, USA) supplemented with vascular endothelial growth factor (200 ng/ml, PeproTech), and forskolin (2 μM, Sigma-Aldrich, USA) (Day 4), and again after 24 h (Day 5). After 24 h, cells were sorted against vascular endothelial cadherin (VE-Cad) (Abcam, United Kingdom) with magnetic assisted cell sorting (MACS) using a Dynamag magnet (Invitrogen, USA) (Day 6). Cells were cultured on fibronectin-coated cell cultures flask in endothelial growth media 2 (Lonza, Switzerland, EGM-2) until use.

### Doxorubicin Supplemented Media (DOX-media) Preparation

Doxorubicin (100 mM) was diluted in DMEM without FBS to create a stock solution of 100 µM, then diluted 1:10 to create a working solution of 10 µM for easy addition of 10 nM. Final concentration necessary was determined according to experimentally determined dosages in literature^127^, and can be compared to *in vivo* measurements using following formula:

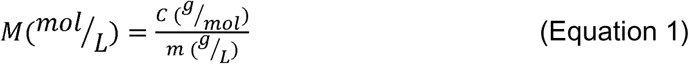

Where m is the molar mass of DOX (543.52 g/mol), C is the blood concentration of DOX (12.54 ng/mL) and M is the desired molarity (10-100 nM, depending on cohort tested).

### Clinical Plasma Sample EV Isolation

Whole blood was collected from patients via direct veinous puncture into ethylenediaminetetraacetic acid (EDTA)-treated tubes to prevent coagulation. Patients were selected for their risk of developing cardiotoxicity, either high-baseline risk or low to moderate risk (n=4) (Supplemental Table S6).To isolate the plasma, whole blood was centrifuged at 1000g 5for minutes at 4°C. Patient plasma was aliquoted to sterile RNAse-free tubes and shipped from the Lambe Institute for Translational Research at the University of Galway to the University of Notre Dame at -80°C. A temperature sensor was included during transport to ensure appropriate storage conditions were maintained. Upon arrival, samples were stored at -80°C until use.

### Extracellular Vesicle (EV) Generation and Isolation

Conditioned media was generated by culturing cells in exosome-free media (no change in CM (+), DMEM Complete and EGM-2 substitute FBS for exosome-free FBS. Conditioned media was centrifuged three times at 500g for 10 min, 2500g for 20 min, and 10,000g for 30 min, and the pellet discarded after each centrifugation step to remove any remaining insoluble matrix remnants. The final supernatant was centrifuged at 100,000g at 4°C for 70 min using an ultracentrifuge (Optima MAX-XP Tabletop Ultracentrifuge, Beckman Coulter). The pellet was either used immediately or stored dry at -80°C. Plasma samples were similarly processed to isolate EVs and pellet was stored accordingly.

### Nanoparticle Tracking Analysis (NTA)

Single pellets were resuspended in 1mL of sterile, particle-free PBS and measured using a NanoSight NS300 machine (Malvern Panalytical) and NTA software version 3.2.16. This method obtains the hemodynamic diameter and concentration of nanoparticles with diameters from 10-1000 nm in solution via Brownian motion analysis. Samples were kept at 4 °C until measurement, and measurements were taken at RT.

### Western Blot

Pellets were lysed in RIPA buffer containing 1% proteinase inhibitor cocktail (Brand, Country) at 4°C for 30 minutes, then protein concentration was assessed via bicinchoninic acid (BCA) assay (Pierce Chemical). Equal amounts of protein were separated by 12% SDS-PAGE and transferred to blotting membranes, which were incubated overnight at 4°C with the rabbit polyclonal primary antibodies against CD9 (Abcam, ab223052), CD63 (Abcam, ab216130), and TSG101 (Abcam, ab30871) at 1:2000 dilution, then for 1 h at RT with HRP-conjugated goat anti-rabbit secondary antibody (Abcam, ab205718). Membranes were then exposed to a chemiluminescent substrate (Clarity ECL, Bio-Rad) and imaged using a ChemiDoc-It2 imager (UVP, Analytik Jena) equipped with VisionWorks software. Images were processed using ImageJ (NIH).

### Ultraviolet-Visible Light (UV-Vis) Spectrophotometry

Pellets were resuspended in 50 µL PBS immediately after ultracentrifugation, and analyzed using a microvolume spectrophotometer (Nanodrop 2000, Thermo Fisher Scientific). For controls, an empty PBS sample, DOX-spiked PBS sample, and supernatant from DOX-cell EV ultracentrifugation we all run as well. Absorbance was quantified at approximately 480 nm, where DOX natively exhibits a broad absorption peak (specifically between 470 nm and 500 nm^128^) (Supplemental Figure S3).

### Doxorubicin-treated cell EV (DOX-EV) Conditioned Media Generation

DOX media was prepared for DOX as described above, with final DOX concentration of 10 nM in media. After 48 h of treatment, media was collected and EVs were isolated. Media conditioned with EVs isolated from DOX-treated cells was created through the addition of isolated EVs to the desired media at 25 µg/mL EVs/media volume. EV mass was measured by the bicinchoninic acid (BCA) assay, as is standard^129^, before addition to media. Before treatment, cells were washed 3x with sterile PBS, then room temperature control or treatment media was added and left undisturbed for 48 h at 37°C, after which cells were again washed 3x with PBS and assays were performed.

### Cardiomyocyte Beating Characterization

To analyze the contractility of iCMs, a block-matching algorithm was performed using MATLAB as described previously^130^. Briefly, iCM or 3D structures were recorded in brightfield in real time under a microscope (Axio Observer. Z1, Zeiss, Hamamatsu C11440 digital camera) for 30 s intervals. Videos were then uploaded to the analysis software, and beating velocity and frequency were calculated, and contraction heat maps were plotted.

### Ca^2+^ Flux Assay

Contraction kinetics were measured by analysis of calcium (Ca^2+^) flux over the tissue, as previously described^131^. Briefly, iCMs were removed from media, which was stored warm and with separated biological replicates, and was incubated with Fluo-4 AM (Thermo Fischer Scientific, USA) according to the manufacturer’s protocol for 30 min at 37°C. After incubation, the staining solution was removed, and the original cell media was returned to the cells. Stained cells were then immediately recorded under a microscope (Axio Observer. Z1, Zeiss) for 30 s intervals, with recording being performed using a green fluorescent channel at 200 ms exposure. Videos were analyzed utilizing an in-house MATLAB code as previously described.

After analysis, resulting waveforms were converted into the frequency domain via 1-dimensional Fast Fourier Transform (FFT) and were subsequently deconstructed into component sine waves, the frequencies of which were plotted.

### Immunostaining

Cells or constructs were washed 3x with PBS to remove residual media, then incubated in 4% paraformaldehyde for 15 min, followed by 0.1% Triton-X for 30 min, then 5% goat serum in PBS for 2 h, all with 3x PBS washes in between. Cells were next incubated with rabbit anti-cleaved caspace-3 (Abcam) and mouse anti-α-SMA (Abcam) primary antibodies (1:100 in 5% goat serum) overnight at 4°C. The cells were then washed and incubated with Alexa Fluor 647-labelled anti-rabbit IgG and Alexa Fluor 488-labelled anti-mouse IgG secondary antibodies (dilution: 1:200 in 5% goat serum) at 4°C for 6 h. Finally, the cells were incubated with DAPI (dilution: 1:1000 in PBS) for 15 minutes at RT and imaged with a fluorescent microscope (Axio Observer. Z1, Zeiss).

### Measurement of Cell Metabolic Activity

DOX media and DOX-EV media were prepared for cells as described above, and cells were treated for 48 h with either DOX media, DOX-EV media, or media with a PBS blank. After 48 h, cells were washed, and media was swapped out for a 10% AlamrBlue (Thermo Fisher)-media mixture^132^. Cells were cultured in the 10% solution for 1 h, after which the solution was transferred to a 96-well plate and the absorbance of each well was measured on a plate reader.

### Reactive Oxygen Species Detection

DOX media and DOX-EV media were prepared for cells as described above, and cells were treated for 48 h with either DOX media, DOX-EV media, or media with a PBS blank. After 48 h, accumulation of reactive oxygen species (ROS) was assessed by a Mitochondrial ROS Assay Kit (eEnzyme, USA). Briefly, the stain solution was prepared as instructed and the cells were incubated at 37℃ for 30 min. Fluorescent images were taken immediately following incubation.

### Seahorse XF ATP Real Time Assay

Seahorse respirometry, to measure oxygen consumption rate (OCR) of cells, was performed with the Seahorse XF Extracellular Flux Analyzer (Agilent, Germany) as reported previously^133^. Briefly, differentiated iCMs, iCFs, and iECs were detached and reseeded into a 96-well Seahorse-compatible plate at 4 × 10L cells/well for iCFs and iECs, and 8 × 10L cells/well for iCMs. iCMs were plated 48 hours earlier to allow additional settling time, which did not affect cell count since iCMs are non-proliferative. iCFs and iECs were seeded and allowed to attach overnight. After attachment, cells were treated for 48 h with DOX media, DOX-EV media, where the EVs came from DOX-treated iCFs or hMFs, or a PBS blank.

Following treatment, cells were stained with Hoechst 33342 (Thermo Scientific, 8 µM) for 30 minutes at 37°C, and cell count in each well were subsequently quantified through ImageJ image analysis. Then, the media was replaced with the provided Seahorse buffer. OCR was then assessed at basal level and following sequential metabolic perturbations with inhibitors prepared at the following final concentrations: 2.5 μM oligomycin, 2 μM FCCP, and 2.5 μM rotenone/antimycin A (Rot/AA). Analysis was performed in the Agilent Seahorse online software, and analyzed data was exported to excel for statistics and graphed in R.

### 3D Cardiac Model Construction

Type 1 collagen (3 mg/ml) (HumaBiologics, AZ, USA) was pH-adjusted to 7 through titration and then refrigerated at 4°C till use. GelMA, synthesized and dissolved in PBS (20% w/v) and maintained in a water bath at 37°C until a fully homogeneous solution was obtained. Then, Irgacure2959 photoinitiator (PI, Sigma, MO, US) stock solution was prepared in PBS (1% w/v) and was added to GelMA. The components were mixed to achieve a bioink with final concentrations of 10% GelMA, 1 mg/ml collagen type 1, and 0.025% PI as previously reported^126,133^. iCMs, DOX-free conditioned media-treated iCFs (control), and pre-treated iCFs were detached using trypsin-EDTA. After centrifugation at 1000 rpm for 5 minutes, iCMs (15 mil/mL) were combined with both types of iCFs (5 mil/mL) and centrifuged again. The supernatant was removed, and each cell pellet was mixed with collagen and GelMA solution, respectfully. Then, bioink was loaded into separate syringes and submerged in the ice for a minute to achieve the required consistency for printing. Droplets were bioprinted using CELLINK BioX6 Bioprinter at a speed of 3 mm/s using 22G nozzles on a sterilized charged glass in a 60- mm dish. Lastly, 3D bioprinted constructs were photo-crosslinked for 30 seconds under UV exposure (6.9 W/cm2) and placed in a well-plate with CM (+) and DMEM complete.

### miRNA Isolation from EVs

RNA was isolated from LVVs and clinical plasma samples using the Total Exosome RNA & Protein Isolation Kit (Thermo Fisher Scientific) using manufacturer’s protocol. Isolated LVVs were resuspended in exosome resuspension buffer and incubated with an equal volume of denaturation solution at 4 °C for 5 min. The solution was then mixed with an equal volume of Acid-Phenol:Chloroform by vortexing for 30 seconds and centrifuged for 5 min at 15,000g. The resulting aqueous phase was extracted and combined with 1.25x volume of 100% ethanol, then transferred to the provided spin column. The spin column was centrifuged at 10,000g for 15 seconds to bind and wash the RNA, then the RNA was eluted in the provided elution solution and quantified via a microvolume spectrophotometer (Nanodrop 2000, Thermo Fisher Scientific).

Following isolation, the eluted miRNA was concentrated using 3 kDa microcentrifuge spin filters (Amicon) according to a previously established protocol^134^. Briefly, the 100 µL miRNA solution was worked up to 420 µL with RNAse-free water and placed into a filter, then centrifuged at 14,000g for 90 minutes. Next, the filter was inverted into a fresh collection tube, and centrifuged at 8,000g for 2 minutes. The resulting isolate is 20-25 µL of concentrated miRNA, which was quantified by the same microvolume spectrophotometer used previously.

### Profiling of Total miRNA Content

Concentrated miRNA was prepared for miRNA profiling (NanoString). The provided miRNA codeset was mixed with the provided hybridization buffer to produce a master mix, and spike-in miRNA controls were prepared at 200 pm. In order, the master mix, concentrated sample miRNA, spike-in miRNA, and provided probes were mixed in a PCR plate and incubated at 65 °C for 16 h. The hybridized solution was then mixed with 15 µL of provided hybridization buffer, for a total volume of 30-35 µL, and added to the provided microfluidic cartridge. The assay was run with the provided protocol for total miRNA analysis, and data was processed and analyzed using the provided software using the recommended settings. Processed data was exported to a .csv spreadsheet for log10 normalization and Z-scoring and subsequent plotting in R, according to the following formula:

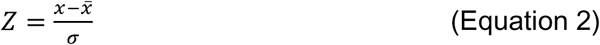

Where z is the Z-score value, x is the log10 normalized count value, x is the average of the log10 normalized count values for a given miRNA, and a is the standard deviation of the log10 normalized count values for a given miRNA.

### miRNA Pathway Analysis

Raw Nanostring data was normalized via Log10 normalization, and uploaded to the Clarivate MetaCore system for pathway analysis. miRNAs were identified by miRBase IDs. Analysis was conducted on the identified target miRNAs to construct a custom network. Automated network analysis was conducted with 50 nodes per network. Results were presented as pathways obtained from the software.

### Statistical Analysis

Results were analyzed by one-way analysis of variance (ANOVA) with post-hoc Tukey’s HSD, two-way ANOVA with post-hoc Tukey’s multiple comparison test, or a two-tailed Student’s t-test with Welch’s correction for unequal standard deviation. Values are presented as the mean ± standard deviation (SD) unless otherwise indicated, and differences were considered significant when p ≤ 0.05.

## ACKNOWLEDGEMENTS

The lyophilization of decellularized ECM was conducted at the Center for Environmental Science and Technology (CEST) at the University of Notre Dame. We thank the Biophysics Instrumentation (BIC) Core Facility for the use of Optima MAX-XP Tabletop Ultracentrifuge. The Nanoparticle Tracking Analysis was conducted using the NanoSight NS300 at the Harper Cancer Research Institute (HCRI) Tissue Core Facility. We thank the Flores-Mireles Lab at the University of Notre Dame for providing access to the NanoString equipment used in this study. The schematics in some figures were created using BioRender.com

## Data Availability Statement

All data required for production of the manuscript is included in this submission. Additional raw data can be provided upon request.

## Ethics Statement

This study was conducted in accordance with the principles of the Declaration of Helsinki. Institutional Review Board approval was not required, as the University of Notre Dame Research Compliance office determined that the work does not constitute human subjects research because the Indiana Donor Network supplies samples without any identifying information.

